# Tape lures swell bycatch on a Mediterranean island harbouring illegal bird trapping

**DOI:** 10.1101/2020.03.13.991034

**Authors:** Matteo Sebastianelli, Georgios Savva, Michaella Moysi, Alexander N. G. Kirschel

## Abstract

Mediterranean islands are critical for migrating birds, providing shelter and sustenance for millions of individuals each year. Humans have long exploited bird migration through hunting and illegal trapping. On the island of Cyprus, trapping birds during their migratory peak is considered a local tradition, but has long been against the law. Illegal bird trapping is a lucrative business, however, with trappers using tape lures that broadcast species’ vocalizations because it is expected to increase numbers of target species. Yet, by how much the use of song playback increases capture rates remains underappreciated. In particular, it is not known whether song playback of target species affects bycatch rates. Here, we show with the use of playback experiments that song playback is highly effective in luring birds towards trapping sites. We found that playback increases six to eight times the number of individuals of target species captured, but also significantly increases bycatch. Our findings thus show that in contrast to popular belief that tape lures are a selective trapping method, they also lead to increased captures of non-target species, which can include species of conservation concern.

## Background

Natural resources are globally threatened due to progressive overharvesting, with animal diversity being particularly affected by its consequences. Birds are very sensitive to anthropogenic impact, which has been a major cause of their decline[2,3]. In addition to indirect impacts on numbers caused by habitat loss and environmental toxification, birds have also been impacted directly, by being targeted for food, the pet trade and sport.

Every year, over two billion birds migrate along the Afro-Palearctic route and concentrate in large numbers around the Mediterranean Basin, which is an important biodiversity hotspot. Mediterranean islands are important stopover sites since they provide trophic resources and shelter for migrant bird species. Humans have long exploited this sudden, seasonal abundance of food resources through hunting. The need to supplement what would in the past have been a low-protein diet has made such habits widespread to the point of making them important culturally in several Mediterranean countries.

The island of Cyprus is an important stopover site for many millions of migrant birds each year comprising over 200 species. Relative to its size, no other country has greater hunting pressure in the Mediterranean basin. Illegal trapping in Cyprus involves killing mostly passerines, and is a common practice well rooted in Cypriot culture. Birds are trapped for food consumption and because of high demand, it is a lucrative business. Eurasian blackcaps (*Sylvia atricapilla*), known locally as ‘ambelopoulia’, are most sought-after by illegal trappers in Cyprus. Though blackcaps are the main target, the use of non-selective trapping methods involving mist nets and lime sticks results in the demise of individuals of many other species. Indeed, of the 155 species recorded captured with mist nets and lime sticks in 2018 in Cyprus, 82 are listed as conservation priority species under the EU Bird Directive or in BirdLife International’s Species of European Conservation Concern, which include the endemic Cyprus warbler *Sylvia melanothorax*.

Capture rates are expected to be amplified by using tape lures: devices involving a loudspeaker to broadcast the songs of target species. Tape lures are usually set by trappers in order to increase catch rates. They are typically played at night to attract nocturnally migrating birds that may hear and respond to the song from great distances, while also reducing detection rates by the authorities.

Use of tape lures to increase catch rates is thought to harm both migratory and resident bird communities, but the extent to which calling devices attract birds to traps has not been quantified. Furthermore, since playbacks are typically aimed at attracting certain target species, we still do not know whether their luring effect is limited to those species or whether they increase the catch rate of other species that may use heterospecific vocalisations as habitat quality cues and for detection of predators.

Here, we aim to quantify with playback experiments the effectiveness of playback of target species’ song stimuli in luring birds into nets. We used recordings of Eurasian blackcap (hereafter blackcap) to determine effects of capture rates of blackcaps and other species compared to controls. We also tested whether playback of Sardinian warbler (*Sylvia melanocephala*) song - a local breeding species’ - also increases capture rates of that species and non-target species. Both species may use vocalizations for conspecific and heterospecific interactions, including in competition for environmental resources. Because of this, we expect a strong response from conspecifics as well as competitor heterospecifics. Through this approach, we aim to determine the extent to which the use of such playback devices increases both catch rates and number of species caught. Our findings would inform authorities and conservation initiatives on the impact on wildlife of the use of tape lures in illicit trapping operations.

## Methods

### Experimental design

We conducted playback experiments between March and October 2016 and in September 2019 at 8 localities in Cyprus (Fig.1). For playback stimuli, we used recordings of blackcap and Sardinian warbler song. Blackcap is a medium sized migrant warbler that does not breed in Cyprus, but occurs in large numbers during migration peaks in spring and autumn, with some individuals overwintering. Sardinian warbler is partially migratory, and found year round in coastal and island populations, including in Cyprus, where it is common and increasing in numbers[30,31].

**FIGURE 1.**
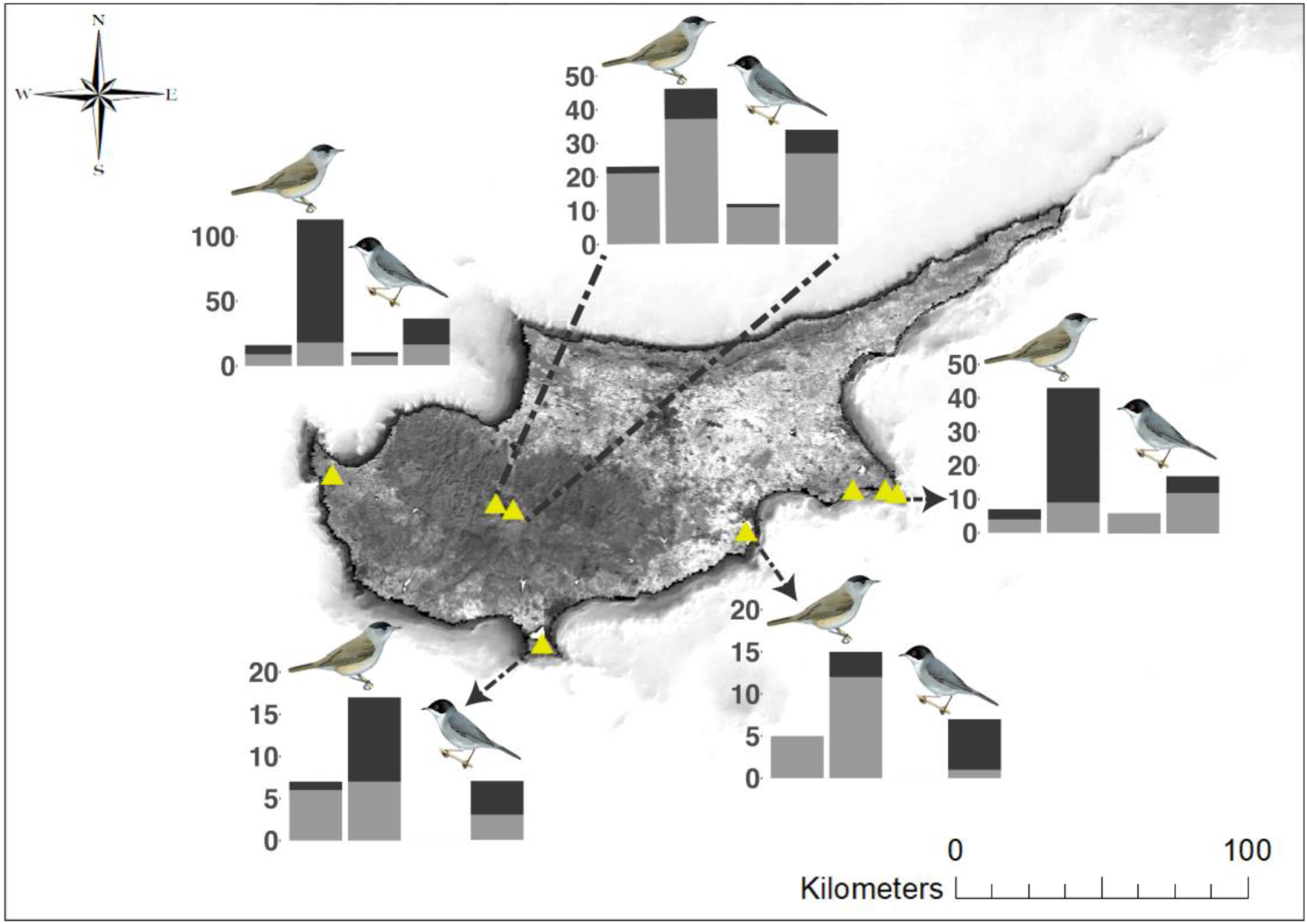
Map illustrating number of heterospecific (grey) and conspecific (black) individuals caught in control (left bar of pair) versus experimental (right bar of pair) nets with blackcap (left pair of bars) and Sardinian warbler (right pair of bars) playback stimuli at different localities: A) Agia Napa and Cape Greco, B) Larnaca Salt Lake, C) Akrotiri Salt Lake, D) Neo Chorio and E) Troodos mountains. Cyprus’ background map illustrates Enhanced Vegetation Index for February 2020. Source: Esri. Bird illustrations courtesy of HBW [27,29].

For each playback experiment session, we used two 12×2.5 m mist nets, with one acting as the experiment (with playback) and the other as control (without playback), and positioned about 100 m away from each other to reduce possible interference between the experiment and control nets. Blackcap and Sardinian warbler stimuli were alternately played in experimental sessions, where a session is an independent experimental period where playback is played continuously for one hour at one of the nets. In all cases, experimental sessions were paired, so that the experimental net in session one was then the control in session two, and vice versa.

Illegal trappers are unlikely to use a large variety of stimuli for a given target species, instead using a single stimulus they have found works well throughout. We aimed to replicate the approach of trappers in using a small number of stimuli in our experiments. We sourced from Xenocanto online repository (www.xeno-canto.org) two blackcap recordings, XC269084 (RMS amplitude = 2712), and XC270439 (RMS amplitude = 2712). For Sardinian warbler we used XC98857 (RMS amplitude = 1180) and our own recording from western Cyprus (RMS amplitude = 1453). Amplitude values were obtained from Raven Pro 1.6.

### Statistical analyses

We first conducted a t-test to determine whether there was an overall effect of playbacks on the total number of captured birds. We then compared Poisson and negative binomial generalized linear mixed models (GLMMs) implemented in R in the lme4 package to investigate how playbacks influence the number of captured birds. Given the slight overdispersion in our data, negative binomial GLMMs provided in all cases the best fit according to the lowest corrected Akaike Information Criterion (AICc) score calculated in the AICcmodavg package in R.

To assess whether playback attracted target species, we ran two models, one with number of captured blackcaps and the other with number of Sardinian warblers as dependent variables. In both models, we included as fixed factors the time, season, and type of playback: a categorical variable with three levels: 1) no playback, 2) blackcap playback and 3) Sardinian warbler playback.

To examine whether playback had an effect on non-target species, we specifically tested the effect of playbacks on all species captured excluding individuals of the species that emitted the given playback stimulus. Specifically, the effect of blackcap playback was tested using as dependent variable the total number of captured birds excluding blackcaps, whereas the effect of Sardinian warbler playback was tested using the total number of trapped birds minus Sardinian warblers. We included time and type of playback as fixed factors. We also included site as a random effect in all our models to account for variation among sites. Models with the best fit (lowest AICc score) for each set of GLMMs were validated by plotting residuals against predicted values and a qq-plot to detect possible deviations from the expected distribution. Model validation functions were provided in the DHARMa R package.

## Results

We caught significantly more birds (t-test: *t=*-4.82; *p =* <0.001) in experimental nets (mean = 2.77, sd = 4.17), catching 333 birds of 31 species, than in controls (mean = 0.75, sd = 1.39), where we trapped 90 birds belonging to 24 species (Fig. 1, Table S1). Numbers of blackcaps were positively affected by both conspecific (GLMM: *z* = 8.33, *p* = <0.001) and Sardinian warbler (*z* = 2.66, *p* = 0.007) playback, with more caught per experiment in spring (*z* = 2.56, *p* = 0.01) compared to autumn (Table 1). By contrast, Sardinian warbler numbers caught were positively affected by conspecific playback (*z* = 5.60, *p* = <0.001) but not blackcap playback (*z* = 1.49, *p* = 0.133), while season also had no effect on their capture rates (Table 1).

**TABLE 1.**
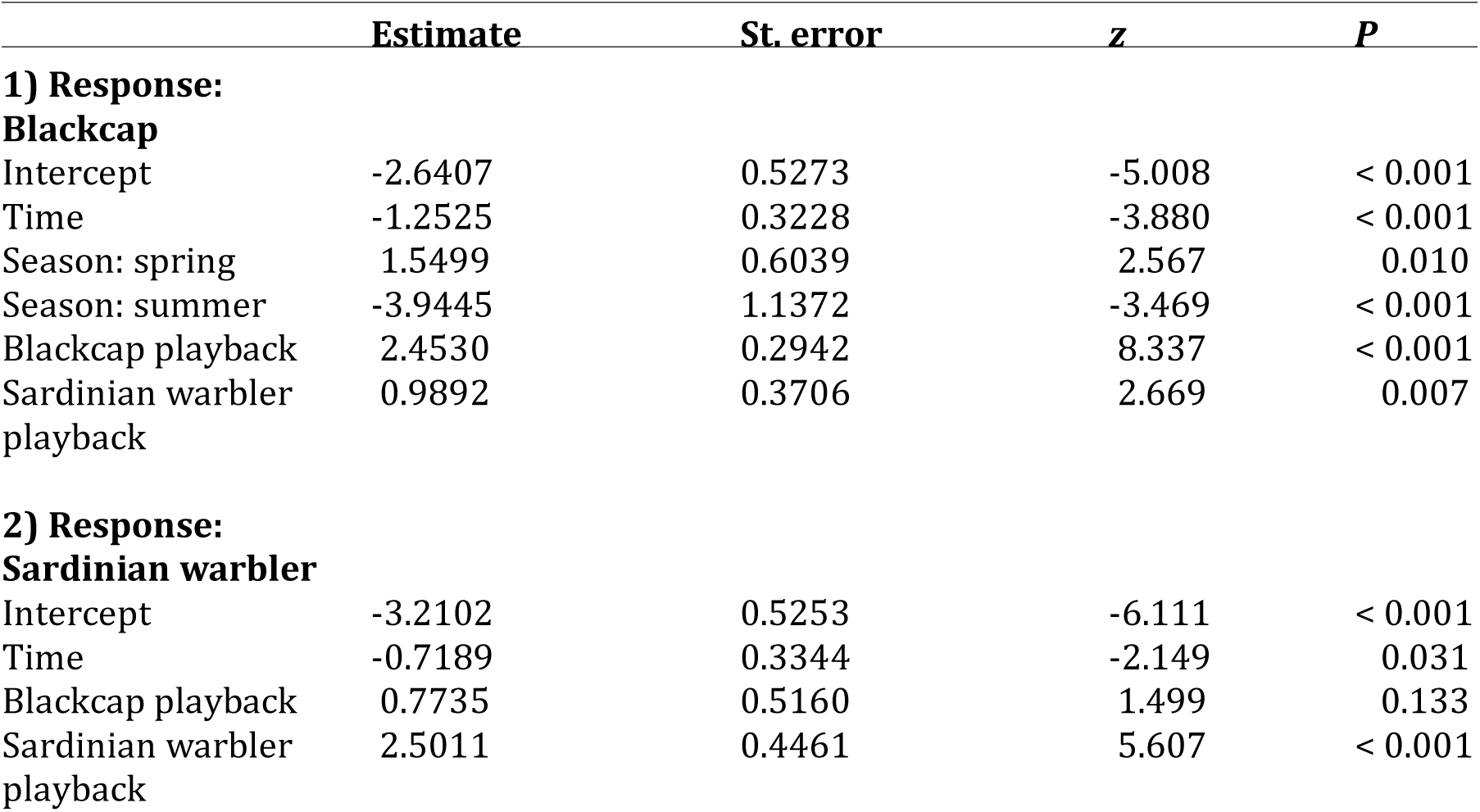
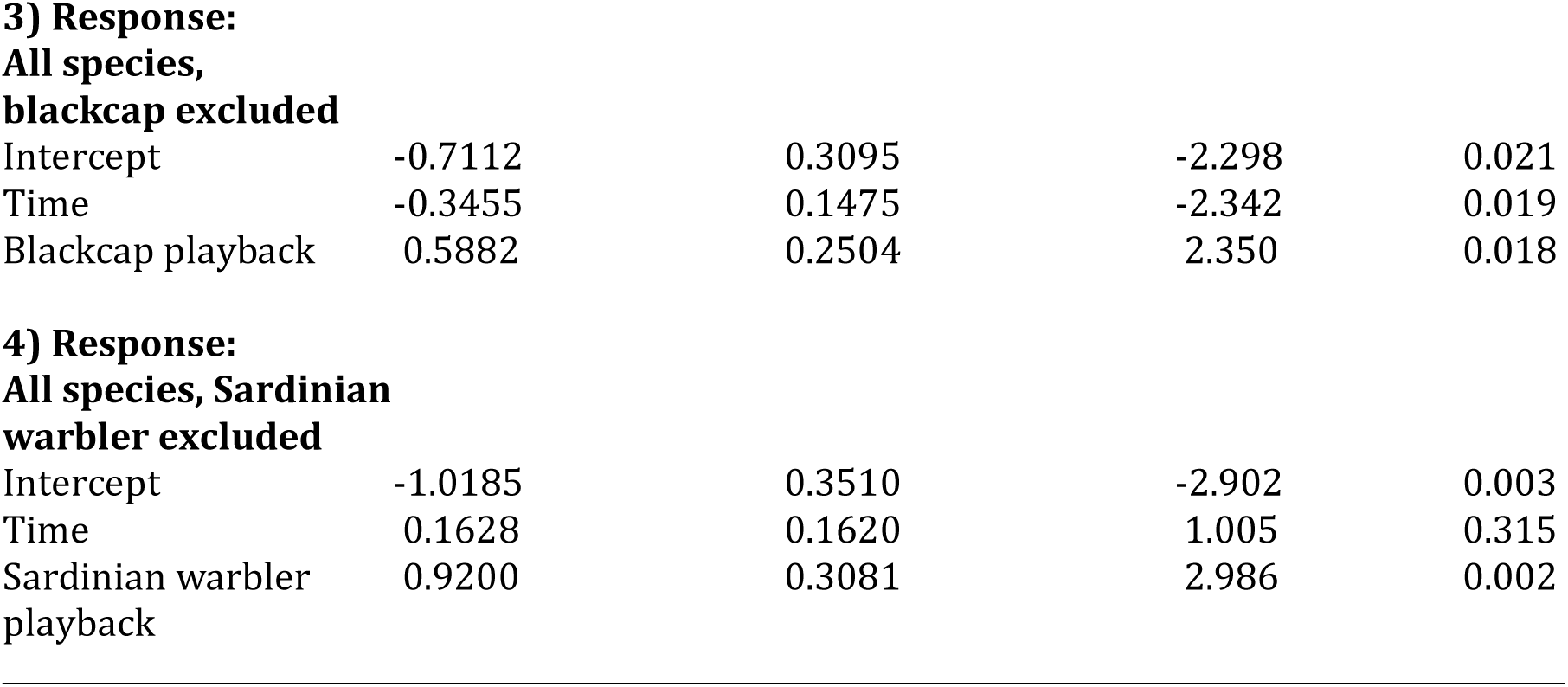
Results of negative binomial GLMMs investigating the effect of 1) blackcap and Sardinian warbler playback on the number of blackcaps caught, 2) blackcap and Sardinian warbler playback on number of Sardinian warblers caught, 3) blackcap playback on heterospecifics minus blackcaps and 4) Sardinian warbler playback on heterospecifics minus Sardinian warblers.

Bycatch numbers were also significantly higher in experimental (n = 106 individuals of 29 species) than in control nets (n = 61 individuals of 22 species) (Fig. 1). Both blackcap (GLMM: *z* = 2.35, *p* = 0.018) and Sardinian warbler playback (*z* =2.98, *p* = 0.002) elicited higher numbers. A significant negative effect of time was also found (*z* = −2.34, *p* = 0.019), but only for bycatch in response to blackcap playback.

## Discussion

Our study shows that the use of tape lures results in an increase from six to eight times capture rates of target species. Tape lures may have a detrimental effect on other avian species since they also attract individuals of non-target species, which would also meet their demise at trapping sites.

Bird song functions in mate choice and territory defence[25,26], so theoretically should only attract conspecific breeding birds, and their close competitors[38]. Indeed, geographic variation in song has been shown to reduce response levels because of differences in dialects[39,40] or resulting from adaptation to habitat differences[41]. Song may also have a function during migration, however, and birds may use conspecific signals to assess environmental quality[42] and trophic resource availability[43]. Season also plays an important role in determining response rates. Spring represents a period of intense migratory activity for many species and this may explain why spring positively affects blackcaps responsiveness in our study. The approach of the breeding season may increase responsiveness as a consequence of increased hormone levels associated with increased territoriality, as demonstrated in other passerines.

According to our results, Sardinian warbler playback also had a positive effect in attracting blackcaps into nets, whereas blackcap playback did not elicit the same response from Sardinian warblers. Blackcaps have been shown to recognize other species[24,28], and the strong response elicited by Sardinian warbler stimuli is likely to have arisen due to similar dietary requirements; both species feed mainly on fruits outside of the breeding season and shift towards an insect-based diet as the breeding season approaches[27,29]. Furthermore, some migrant species use resident species as indicators of habitat-quality and food availability[45] as well as to infer the presence of predators from the calls of resident prey species[46]. We suggest blackcap responses to Sardinian warbler playback also reflect migrants eavesdropping on heterospecific vocalisations for their own benefit. However, because Sardinian warblers are resident species, they are less likely to rely on other species to locate food resources and would not be expected to recognize vocalisations of a species that does not breed in Cyprus, such as blackcap. Notwithstanding, no response by Sardinian warbler to blackcap song might even reflect avoidance behavior, whereby resident species avoid migratory birds to escape competition for food, which has been shown in Sardinian warbler[47]. However, we did not find that Sardinian warbler avoided blackcap experimental nets more than controls.

In our study, we also show that other species responded positively to both blackcap and Sardinian warbler playbacks. Related heterospecific birds such as *Sylvia* warblers (e.g. *S. curruca* and *S. melanothorax*) responded to the calls, possibly because of overlapping diet and habitat requirements[28], thus contributing to the strong positive response to both playbacks. Also, response to heterospecific vocalization might be directly related to phylogenetic relatedness since they tend to share similar song features, as demonstrated in other taxa[48]. Heterospecific responses of more distantly-related species such as willow warbler (*Phylloscopus trochilus*), spotted flycatcher (M*uscicapa striata*), European chaffinch (*Fringilla coelebs*) and common redstart (*Phoenicurus phoenicurus*) may also be elicited by a food expectancy or the advantages of safety in numbers and increased risk detection. Under the latter scenario, eavesdropping heterospecific signals such as warning signals may lead to important advantages such as a rapid response to threats and therefore an optimization of foraging[23].

In this paper, we show that tape lures boost capture rates at trapping sites. The joint use of non-selective traps with tape lures increases the number of both individuals and range of species caught, which often include threatened species or local endemics such as the Cyprus warbler, which might already be in population decline because of other factors such as habitat disturbance and competition with recent colonizers[30]. The tradition of using lime sticks to catch migrant birds for a meal is illegal, much because of the non-selectivity of this methods and because the extensive use of these methods leads to a mass killing of birds in the Mediterranean[15,16,49]. The industrial level illicit trapping of millions of birds using playback devices to lure birds into vast mist nets needs immediate action by the authorities and the continued attention of conservation practitioners. Targeting the source of song playbacks is likely to be the most effective way of finding the traps – it is what attracts the birds into the traps in the first place.

## Acknowledgments

We thank Chrystalla Costi, Sifiso Lukhele, and Emmanuel Nwankwo for assistance in the field, and Martin Hellicar and Stavros Christodoulides for helpful comments on the manuscript. This research was supported by A. G. Leventis Foundation grants to MS and MM.

