## Supplemental Table 1 for "Tape lures swell bycatch on a Mediterranean island harbouring illegal bird trapping"

**TABLE S1.** Number of individual birds captured per each species in the experiment vs. control nets according to the type of playback.

| Species | Total | Experiment | | Type of playback | |
|---|---|---|---|---|---|
| | | Playback | Control | Blackcap | Sardinian warbler |
| *C. chloris* | 3 | 1 | 2 | 1 | 0 |
| *C. cetti* | 1 | 1 | 0 | 1 | 0 |
| *E. calandra* | 1 | 1 | 0 | 0 | 1 |
| *F. coelebs* | 32 | 19 | 13 | 7 | 12 |
| *O. cypriaca* | 4 | 3 | 1 | 3 | 0 |
| *C. brachydactila* | 3 | 1 | 2 | 1 | 0 |
| *F. hypoleuca* | 1 | 1 | 0 | 0 | 1 |
| *L. megarhynchos* | 3 | 1 | 2 | 1 | 0 |
| *L. svecica* | 1 | 1 | 0 | 0 | 1 |
| *M. striata* | 19 | 12 | 7 | 12 | 0 |
| *P. ater* | 6 | 5 | 1 | 3 | 2 |
| *P. major* | 6 | 2 | 4 | 1 | 1 |
| *P. hispaniolensis* | 2 | 0 | 2 | 0 | 0 |
| *P. domesticus* | 5 | 3 | 2 | 1 | 2 |
| *T. merula* | 3 | 3 | 0 | 3 | 0 |
| *P.phoenicurus* | 5 | 4 | 1 | 3 | 1 |
| *P. trochilus* | 22 | 15 | 7 | 14 | 1 |
| *S. atricapilla* | 196 | 175 | 21 | 151 | 24 |
| *S. communis* | 2 | 1 | 1 | 1 | 0 |
| *S. curruca* | 7 | 6 | 1 | 4 | 2 |
| *S. melanocephala* | 60 | 52 | 8 | 10 | 42 |
| *S. melanothorax* | 4 | 3 | 1 | 0 | 3 |
| *S. borin* | 1 | 1 | 0 | 1 | 0 |
| *S. crassirostris* | 2 | 2 | 0 | 0 | 2 |

| | | | | | |
|---|---|---|---|---|---|
| *A. scirpaceus* | 11 | 10 | 1 | 10 | 0 |
| *A. arundinaceus* | 1 | 1 | 0 | 1 | 0 |
| *H. pallida* | 4 | 3 | 1 | 1 | 2 |
| *F. parva* | 1 | 1 | 0 | 0 | 1 |
| *J. torquilla* | 2 | 1 | 1 | 0 | 1 |
| *L. collurio* | 1 | 0 | 1 | 0 | 0 |
| *C. carduelis* | 2 | 1 | 1 | 1 | 0 |
| *S. serinus* | 1 | 0 | 1 | 0 | 0 |
| *G. glandarius* | 1 | 1 | 0 | 0 | 1 |
| *U. epops* | 1 | 0 | 1 | 0 | 0 |
| *T. troglodytes* | 2 | 2 | 0 | 2 | 0 |
